# Effects of transcranial alternating-current stimulation to secondary motor areas on cortical oscillations in stroke patients

**DOI:** 10.1101/529818

**Authors:** Lutz A. Krawinkel, Marlene Bönstrup, Jan F. Feldheim, Robert Schulz, Winifried Backhaus, Till R. Schneider, Jonas Misselhorn, Bastian Cheng, Christian Gerloff

## Abstract

**Background:** There is growing evidence that secondary motor areas are relevant for recovery after motor stroke. Communication among brain areas occurs via synchronization of oscillatory activity which can potentially be modulated via transcranial alternating-current stimulation (tACS).

**Hypothesis:** We hypothesized that tACS to secondary motor areas of the stroke-lesioned hemisphere leads to modulation of task-related connectivity among primary and secondary motor areas, reflected in metrics of EEG coupling in the frequency domain.

**Methods:** We applied focal tACS at 1mA peak-to-peak intensity to ipsilesional ventral premotor cortex (PMv) and supplementary motor area (SMA) in chronic stroke patients while they moved their impaired hand. To probe effects of stimulation on cortical oscillations, several task-related EEG-based connectivity metrics (coherence, imaginary coherence, phase-locking value, mutual information) were assessed before and after each stimulation.

**Results:** Overall, we found significant but weak modulations of the motor network by tACS. Stimulation of PMv reduced task-related coupling between (i) both primary motor cortices (M1) (coherence, −0.0514±0.0665 (mean±SD, active stimulation) vs. 0.0085±0.0888 (sham), p=0.0029) and (ii) between ipsilesional M1 and contralesional PMv (coherence, - 0.0386±0.0703 vs. 0.0226±0.0694, p=0.0283; phase-locking value, −0.0363±0.0581 vs. 0.0036±0.0497, p=0.0097) compared with sham stimulation.

**Conclusions:** In this exploratory analysis, tACS to the ipsilesional PMv induced a weak decrease of task-related connectivity between ipsilesional M1 and contralesional M1 and PMv. As an excess of interhemispheric coupling is under discussion as maladaptive phenomenon of motor reorganization after stroke (e.g., bimodal balance-recovery model), tACS-induced reduction of coupling might be an interesting approach to assist re-normalization of the post-stroke motor network.

## 1. Introduction

Stroke is one of leading causes for disability and especially dysfunction of hand control hampers reintegration into everyday activities (Rathore et al., 2002; Survival et al., 2001). There is growing evidence that secondary motor areas play a major role in the process of lesion-induced reconfiguration of motor network architecture. After a motor stroke, a reinstatement of coupling patterns among the ventral premotor cortex (PMv), the supplementary motor area (SMA) and the primary motor cortex (M1) in the ipsilesional hemisphere appears to be part of the recovery process (Rehme et al., 2011). Particularly coherence of oscillatory activity was shown to be reduced among cortical motor regions in the stroke lesioned hemisphere (Gerloff et al., 2006). Together, these results may indicate a relevance of strong interregional cross-talk among primary and secondary motor areas in the ipsilesional hemisphere for near-healthy motor output.

Activity in the contralesional hemisphere has been ambiguously discussed as either playing a compensatory role with replacement or support of the ipsilesional hemisphere (Johansen-Berg et al., 2002; Lotze et al., 2006). On the other side there are indications that activity in contralesional M1 may be counteracting motor function of the affected arm (Grefkes et al., 2010; Nowak et al., 2008). This ambiguous role has been reconciled in the bimodal balance-recovery model (Di Pino et al., 2014) embracing amount of structural damage as well as recovery stage. Despite the unclear functional role of the contralesional hemisphere, it is certain that contralesional motor areas are behaviorally relevant nodes in the post-stroke network.

To date, there are multiple efforts to enhance recovery, especially by interfering with local activity in strategic network nodes by non-invasive brain stimulation (Hummel et al., 2005; Rossi et al., 2013). However, non-invasive brain stimulation has the potential to modulate both local activity and inter-regional connectivity (Ahn et al., 2019; Ahn et al., 2018; Helfrich et al., 2014; Plewnia et al., 2008; Schwab et al., 2018; Thut et al., 2011a; Veniero et al., 2015; Vernet et al., 2013; Vossen et al., 2015; Zaehle et al., 2010). Particularly the application of tACS is a rapidly evolving field with documented effects on both oscillatory activity and human behaviour (Ahn et al., 2019, 2018; Brinkman et al., 2016; Helfrich et al., 2014; Lustenberger et al., 2016), inducing lasting after-effects (Veniero et al., 2015; Vossen et al., 2015; Zaehle et al., 2010).

Application of tACS effects in stroke patients is sparse but promising. Naros et al. were able to improve neurofeedback-mediated therapeutic intervention with beta-band tACS applied over the stroke lesion and accredited this effect to a variance reduction of brain oscillations immediately following stimulation bouts (Naros & Gharabaghi, 2017).

On the basis of (i) previously identified relevant nodes and connectivity patterns in motor control in healthy and stroke patients (Bönstrup et al., 2015; Gerloff et al., 2006; Rehme et al. 2011) and (ii) the potential of tACS to effect oscillatory activity (Ahn et al., 2019; Helfrich et al., 2014; Schwab et al., 2018), we applied focal tACS to secondary motor areas in chronic stroke patients. We tested network effects of two combinations of stimulation region and frequency, SMA at 21Hz and PMv at 11Hz, guided by earlier works indicating pronounced frequency coupling between both regions and M1 at the respective frequencies (Boenstrup et al., 2014; Bönstrup et al., 2015). Network-effects were investigated by recording neuroelectric activity non-invasively using EEG and calculating task-related modulations of local activity and inter-regional connectivity. Activity was exploratory assessed by means of spectral power and inter-regional connectivity by means of coherence, imaginary coherence and phase-locking value as well as cross-frequency coupling (mutual information).

Our aim was to investigate if focal rhythmic stimulation can modulate task-related connectivity in a lesioned motor network. We opted to directly study this in stroke survivors (i) because the altered cortical oscillation patterns in patients might react differently from those of healthy subjects and (ii) because feasibility of this type of ‘adjuvant’ non-invasive brain stimulation might be different in healthy subjects and patients (and the latter are the actual target population).

## 2. Design, patients and methods

### 2.1. Study design

Patients underwent three blocks of transcranial alternating current stimulation (tACS) in one experimental session (Fig. 1): (i) 11Hz tACS was applied over ipsilesional PMv, (ii) 21Hz tACS over SMA, (iii) sham stimulation in a balanced manner over PMv (11Hz) or over SMA (21Hz). The order of stimulation blocks was pseudo-randomized to avoid order effects. To additionally minimize carry-over effects, the sham stimulation block was always placed between the two target stimulation blocks (i.e., PMv-sham-SMA or SMA-sham-PMv). During each stimulation block, patients performed visually instructed and guided grip force production tasks with the affected hand (20 trials). Following each stimulation block, the task was repeated in 20 trials without any stimulation. These blocks were included to measure outlasting stimulation effects on task-related brain rhythmic activity.

**Figure 1.**
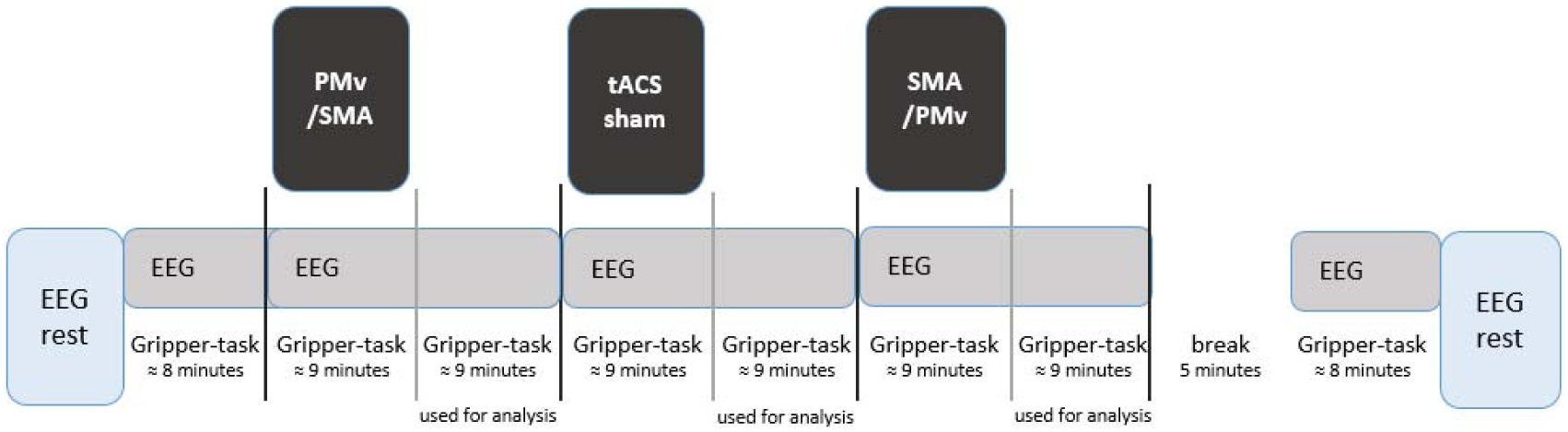
After each stimulation an “offline”-block without tACS stimulation was performed. “Offline”-blocks were used for analysis of EEG data. Before and after stimulation blocks, patients underwent a resting-state EEG and performed a behavioral baseline.

At the beginning and the end of the session, patients (i) underwent a resting-state EEG (4 minutes) and (ii) performed the task as a pre- and post-task-baseline with a five-minute break before the post-baseline. Thus, in total the task was repeated eight times (8 × 20 trials, Fig. 1). The session ended with a thorough neurological exam and standardized testing (Nine Hole Peg Test, measurement of grip force, Upper Extremity Fugl-Meyer-Score (UEFM), National Institute of Health Stroke Scale (NIHSS)).

### 2.2. Patients and clinical testing

16 chronic stroke patients (Tab. 1, for lesion overlay see Fig. 2) were recruited for this study. Some of these patients had already participated in previous experiments (Bönstrup et al., 2018; Schulz et al., 2016). The interval after stroke event was at least one year and a half. Patients had a mean age of 62.3 years (±10.1 (SD)), mean interval after stroke was 35.3 months (±11.0). Our cohort included 4 female and 12 male patients. Patients were generally well-recovered with a mean UEFM of 63.0 (±12.0) and a NIHSS of 0.8 (±1.5). Initial NIHSS was 4.4 (±2.3) and initial UEFM was 47.9 (±18.2). Contraindications against non-invasive brain stimulation were checked by a safety questionnaire (Rossi et al., 2009). All patients gave written informed consent according to the declaration of Helsinki. The study has been approved by the local ethical committee of the Medical Association of Hamburg (PV4931).

**Table 1.** Clinical characteristics of patients. Clinical data. M indicates male, F female. Age is given in years. BG indicates basal ganglia; CI, internal capsule; CR, corona radiata; INS, insula; ITG, inferior temporal gyrus; LOC, lateral occipital cortex; MED, medulla oblongata; MFG, middle frontal gyrus; MTG, middle temporal gyrus; PON, pons; POC, postcentral gyrus; PRC, precentral gyrus; SFG, superior frontal gyrus; SPL, superior parietal lobule; STG, superior temporal gyrus.

| Sex | Age | Handedness | Stroke / Side of Stimulation | Stroke Location | Initial NIHSS | Initial UEFM | Months after Stroke | Residual NIHSS | Residual UEFM |
| --- | --- | --- | --- | --- | --- | --- | --- | --- | --- |
| M | 66 | right | left | CI,CR | 3 | 37 | 52 | 0 | 66 |
| M | 72 | left | right | CI,CR,BG,INS | 3 | 62 | 56 | 1 | 66 |
| F | 71 | right | right | PON | 5 | 61 | 45 | 0 | 66 |
| M | 57 | right | left | MED | 4 | 62 | 45 | 0 | 66 |
| M | 53 | right | right | CI,BG,INS,MFG | 7 | 20 | 42 | 0 | 66 |
| M | 55 | right | right | CI,BG | 4 | 65 | 41 | 0 | 66 |
| F | 58 | right | right | PRC | 1 | 63 | 35 | 0 | 66 |
| M | 51 | right | left | CR,SPL,MFG | 3 | 56 | 30 | 0 | 66 |
| M | 66 | right | left | CR | 3 | 42 | 39 | 0 | 66 |
| F | 72 | right | right | CR,BG,INS | 5 | 56 | 29 | 0 | 66 |
| M | 67 | right | right | CI, CR | 8 | 6 | 38 | 6 | 18 |
| M | 46 | right | left | CR,PRC,MFG,SFG,POC,LOC,MTG,ITG,STG | 8 | 61 | 26 | 1 | 66 |
| F | 80 | right | left | CI,BG | 0 | 66 | 25 | 0 | 66 |
| M | 55 | right | left | BG,CI | 5 | 41 | 21 | 1 | 66 |
| M | 77 | right | left | PONS | 5 | 26 | 17 | 1 | 66 |
| M | 50 | right | left | BG, CI CR, STG,INS | 7 | 43 | 24 | 2 | 66 |

**Figure 2.**
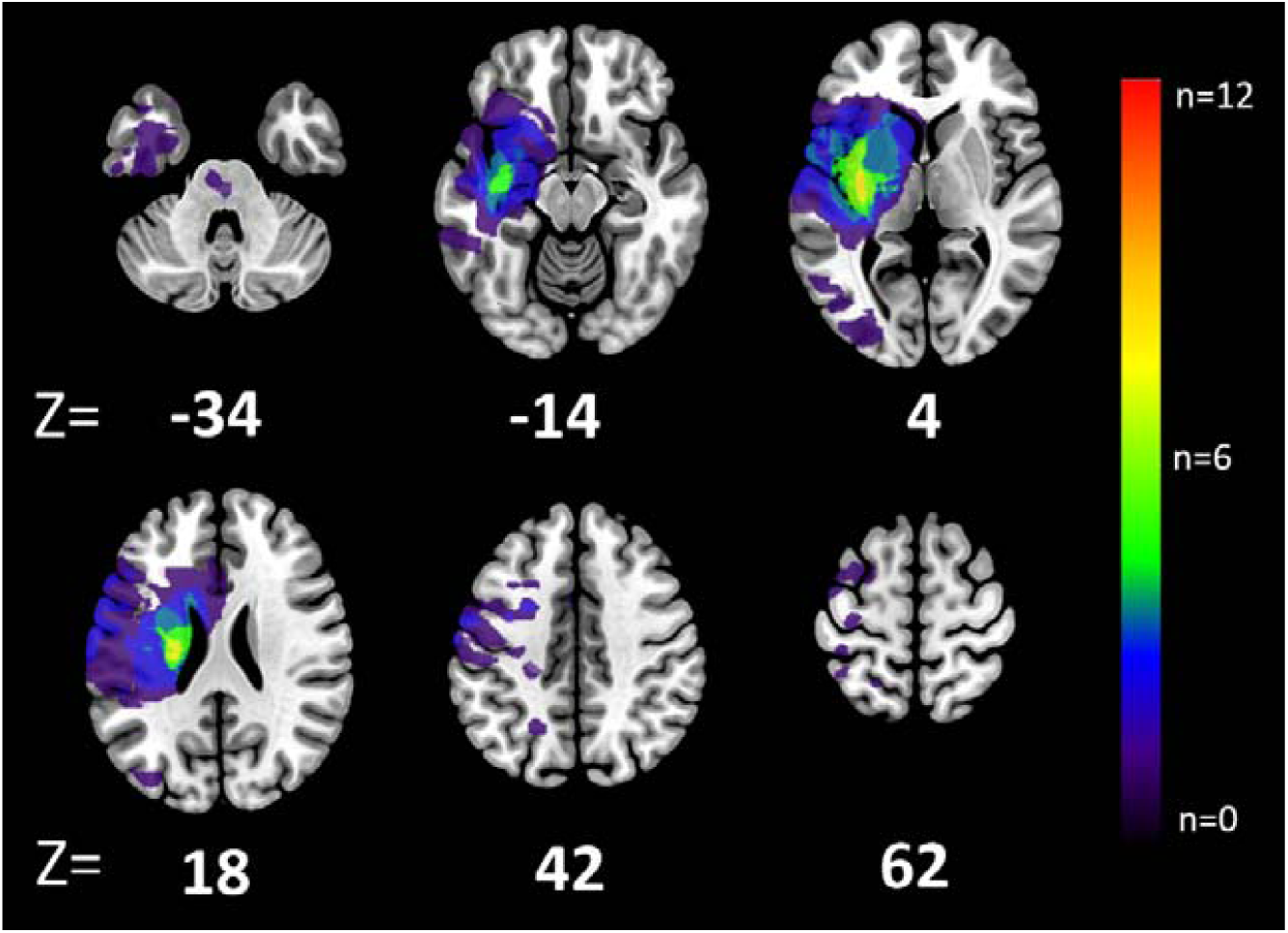
Lesion overlay. Summary overview of lesion locations in all 16 stroke patients with color indicating frequency of lesions. For illustrative purposes, all lesions were mirrored to one hemisphere. All lesions were segmented on Fluid-attenuated inversion recovery (FLAIR) images on days 3-5 after stroke and consecutively registered to Montreal Neurological Institutes (MNI) normal space. Overlays are displayed on a T1 template in standard space, z-values of representative slices are given.

### 2.3. Motor paradigm

We used a motor task well-established in stroke patient cohorts (Bönstrup et al., 2014; Bönstrup et al., 2018; Bönstrup et al., 2016). It required the patients to perform simple, isometric whole-hand grips with the affected hand. The motor task consisted of 20 visually guided whole-hand grips in an event-related design with an inter-trial interval of 12±2s and a stable, isometric hold-phase of 9s. Applied force was equaled 20% of the maximal voluntary contraction (Fig. 3).

**Figure 3.**
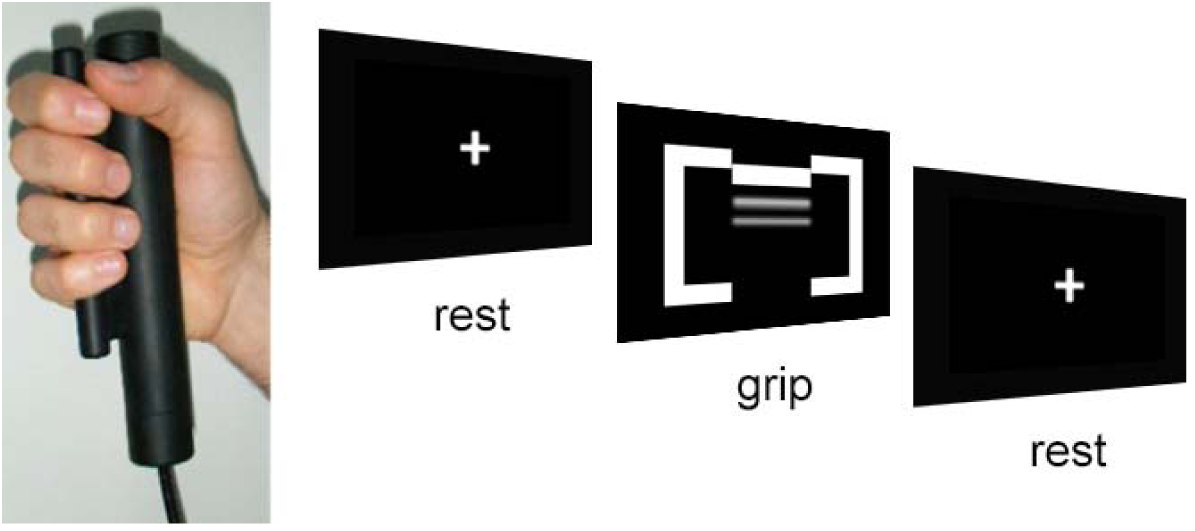
At the presentation of two opposing brackets flanking a horizontal bar, patients had to perform a grip with their affected hand. This resulted in lifting of the bar towards the upper limit of the brackets (target). The required force to reach the target equaled 20% of maximum voluntary contraction. After 9s of holding this force level, the brackets and bar disappeared and a fixation cross reappeared.

### 2.4. Transcranial alternating current stimulation

TACS was applied by a battery-operated stimulator (DC-Stimulator Plus, NeuroConn, Ilmenau) using five Ag/AgCl electrodes (12mm diameter, mounted in a circular arrangement of four outer electrodes and a central electrode, Easycap). The combined impedance of all stimulation electrodes was kept below 5kΩ, as measured by the NeuroConn device. Single impedances of stimulation electrodes being below 5kΩ where additionally measured by an independent impedance meter (SIGGI II, BrainProducts, Gilching, Germany). For PMv, the affected hemisphere was stimulated. For SMA, stimulation was applied over the midline without attempt to differentiate between left and right SMA, as this was beyond the spatial resolution of the tACS montage. For PMv stimulation, FC5 (FC6) was the central electrode, FFC5, FCC5, FT7h and FC5h (FFC6, FCC6, FT8h and FC6h) were the outer electrodes. For the SMA stimulation, FCz was the central electrode, FFCz, FCCz, FC1h and FC2h were the outer electrodes (Fig. 4). A sinusoidal alternating current of 1mA (peak-to-peak (Helfrich, Knepper, et al., 2014; Helfrich, Schneider, et al., 2014)), was applied continuously for 8 minutes over PMv (11Hz) stimulation and over SMA (21Hz). The choice of these particular frequencies was based on earlier work indicating pronounced frequency coupling between the target regions and M1 at these frequencies (Boenstrup et al., 2014, 2015). The current was ramped up over 4s to 1mA, but was ramped down over 4s after 10s in the sham condition. In 8 patients, sham stimulation was applied over PMv (11Hz), in another 8 patients sham stimulation was applied over SMA (21Hz). Like the order of verum stimulation, the two sham-stimulation conditions were randomized across patients. TACS was applied during the trial and inter-trial baselines for all 20 trials of the motor task and then faded out.

**Figure 4.**
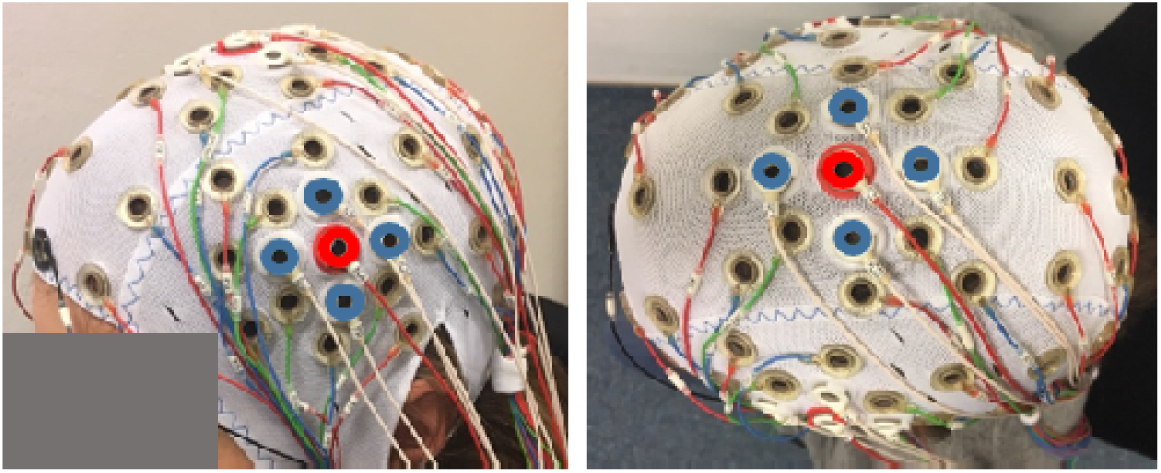
Montage for tACS stimulation of PMv and SMA. For ipsilesional PMV stimulation, FC5 (FC6 respectively, red) was the central electrode (left panel). FFC5, FCC5, FT7h and FC5h (FFC6, FCC6, FT8h and FC6h respectively, blue) were the outer electrodes. For SMA stimulation (right panel), FCz was the central electrode (red) while FFCz, FCCz, FC1h and FC2h (blue) were the peripheral electrodes (right panel).

### 2.5. EEG

#### 2.5.1. Data acquisition

Continuous EEG was recorded from 63 Ag/AgCl electrodes arranged in the 10/20 system (easyCAP®, Brain Products GmbH, Gilching, Germany). We included additional electrode positions close the stimulation-sites as FFT7h/FFT8h, FTT7h/FTT8h, FFC5h/FFC6h, FCC5h/FCC6h, FFC1h/FFC2h and FCC1h/FCC2h. Impedances were kept below 10kOhm. Data were sampled at 5000Hz, amplified in the range of ±16.384mV (resolution 0.5µV). Recordings were referenced to a nose-tip electrode during recording (BrainAmp MR Plus® amplifier, Brain Products GmbH, Gilching). One electrode was mounted below the left eye for Electrooculogram. Electrode positions were registered using an ultrasound localization system (CMS20, Zebris, Isny, Germany) before the EEG-recording. EMG activity of the left and right flexor and extensor digitorum was recorded using a bipolar montage with a tendon reference near the wrist (brainAmpExG, Brain Products GmbH, Gilching). Trial begin and end were automatically documented with markers in the continuous recording.

#### 2.5.2. Data analysis

Data analysis was performed with the FieldTrip package for EEG/MEG data analysis (Oostenveld, Fries, Maris, & Schoffelen, 2011), the GCMI (https://github.com/robince/gcmi) toolbox, the MEG&EEG toolbox of Hamburg (https://www.uke.de/dateien/institute/neurophysiologie-und-pathophysiologie/downloads/meth.zip) and SPM12b (University College London, UK) on MATLAB 17a (The Mathworks Inc., Massachusetts, USA).

##### Preprocessing

The continuous EEG was offline down sampled to 125Hz, detrended, notch-filtered and subjected to an independent component analysis (logistic infomax ICA; (Makeig et al., 1996)) to remove eyeblink artifacts. The 20 trials of each condition were then segmented in epochs of 1s duration comprising the 4s preceding each task onset for an in-trial baseline and covering the isometric hold phase, starting 1s after the beginning of each trial. Trials were then visually inspected to reject remaining artifacts (number of 1s long trials after artefact rejection (mean±SD): task 119±14, rest 235±29).

##### Source reconstruction

Segmented data were re-referenced to a common cephalic average and band-pass filtered in the frequency range of interest (11±1Hz, 21±1Hz) corresponding to the stimulation frequencies using the default filter settings of the FieldTrip toolbox. For the connectivity analysis, we reconstructed source space activity and connectivity at a defined network comprising bilateral PMv and M1 and SMA. The coordinates were obtained from a previous fMRI experiment (Schulz et al., 2016), the coordinate of the SMA was averaged due to the close proximity of both SMA that exceeds the spatial resolution of the EEG (Nunez et al., 2015). For power analysis, we reconstructed source space activity on a regularly spaced (14mm) three-dimensional grid of locations within the brain volume (AAL atlas, provided by the FieldTrip package). Individual forward models were computed as described in an earlier report (Bönstrup et al., 2018).

##### Spectral Power and Connectivity Analysis

To obtain whole brain spectral power estimates, each source space time series on the cortical grid was Hilbert transformed and the power at each time point was derived as the absolute value of the Hilbert transformed time series.

Brain regions can functionally connect by synchronization of their oscillatory neural activity which can be quantified by coherence, i.e., correlation of rhythmic signals including phase and amplitude, phase-locking value, i.e. emphasizing phase synchrony between nodes (Bastos & Schoffelen, 2016) or imaginary part of the coherence, being robust to spurious coherence arising from electric field spread which mainly manifests as phase difference of 0° or 180° (Bastos & Schoffelen, 2016; Nolte et al., 2004). This wide analytic approach was chosen to increase robustness of results. Moreover these metrics were used for evaluating effects of tACS in previous studies (Chander et al., 2016; Helfrich, Knepper, et al., 2014; Schwab et al., 2018).

For each metric, the source-space time series were first Hilbert-transformed. Cross-spectra between two locations’ time series were calculated as the mean of the product of the analytic signal of one and the conjugate of the analytic signal of the other time series over all samples. Then, coherence values were calculated as the cross-spectra divided by the product of the corresponding auto-spectra according to the formula

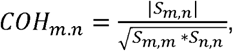

where S denotes the cross-spectrum at any given frequency, averaged over samples, and m and n are nodes. For the analysis of the phase-locking value between each node, the phase signal of each nodes’ time series was first derived as the analytic signal divided by the absolute of the analytic signal. The phase-locking value was then computed as the mean of the product of the phase signal of one and the conjugate of the analytic signal of the other time series over all samples, according to the formula:

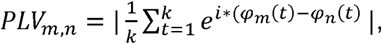

where *φ* denotes the signal phase at any given frequency, *t* the sample number and k the total number of samples.

To obtain connectivity estimates between the activity of the nodes of the motor network across the stimulation frequencies, we calculated the non-linear mutual information between the power envelopes of each node at each frequency combination with parametric Gaussian copula transformation (function gcmi_cc with default settings (Ince et al., 2017)).

For normalization of the underlying distribution, power estimates were log-transformed and connectivity estimates were subjected to a hyperbolic inverse tangent (tanh-1) transformation (Bönstrup et al., 2018; Gerloff et al., 1998, 2006). As a second step, to reduce inter-subject variability, the estimates recorded during task execution were normalized with the estimates during in-trial baseline via subtraction (Bönstrup et al., 2018; Gerloff et al., 1998, 2006). Task-related power decrease (TRPD) thereby reflects ‘activation’ (Gerloff et al., 2006; Hummel & Gerloff, 2006).

##### Statistical Testing

Statistical tests were done using the LMER package in R (http://www.r-project.org/). Linear mixed-effects models were calculated for each connection, or regional activity, and stimulation condition (PMv, SMA) with fixed effects condition (stimulation, sham), as the main effect of interest, and the order of stimulation (sham or verum stimulation first). The patient ID was modelled as a random intercept to account for individual adjustments of the target effect. Each effects’ predictive power was assessed using analysis of variance and a level of p<0.05 determined as significant.

In general, the present study was designed as an explorative analysis. Therefore, we primarily decided to report uncorrected P-values. However, to allow the reader a judgment of the statistical significance of the individual results, critical P-values after correction for multiple testing (Benjamini & Hochberg, 1995) are given in the Supplementary Information.

##### Subject exclusion

Due to excessive muscle artifacts, we excluded 2 patients for alpha-band and 6 patients for beta-band analysis. After exclusions, the alpha-band analysis included 8 patients with randomization order ‘PMv-sham-SMA’, 6 patients with ‘SMA-sham-PMv’, the beta-band-analysis (and cross frequency-analysis) was based on 6 patients with the order ‘PMv-sham-SMA’, and 4 patients with the order ‘SMA-sham-PMv’.

### 2.6. Simulation of electric fields

We computed estimates for strength and focality of the electric fields resulting from tACS. To that end, a leadfield matrix *L* was computed using a three-shell head model and a cortical grid in MNI space (Nolte & Dassios, 2005). We calculated the electric field 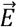 at location 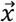 as the linear superposition of evoked fields of all injected currents *α*_*i*_ at stimulation electrodes *i* = {1,2, …,5} as

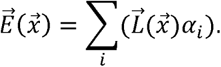

Thereby, we obtained a cortical distribution of electric field strengths.

## 3. Results

### 3.1. Feasibility in stroke patients

According to the patients, stimulation was mainly noticeable as a weak tingling during the ramp-in phase. Patients did not report any unspecific side effects. No severe side effects like seizures occurred during or after stimulation.

### 3.2. Task-Related-Power (TRPow)

Local activity was quantified as spectral-power changes (section 2.5.2.) at each region within the motor network comprising ipsilesional and contralesional PMv, SMA, ipsilesional and contralesional M1. These nodes and their connections have been shown to be activated during whole-hand grip, especially their activation patterns differed from healthy controls (Bönstrup et al., 2015, 2016; Rehme et al., 2012). Figure 5A shows TRPow, averaged over all conditions for the alpha- and beta-band, reproducing published activation patterns (Bönstrup et al., 2016). Task-related power decrease (TRPD) during motor execution (desynchronization) was most prominent at both primary cortices for the alpha band and at ipsilesional M1 (contralateral to the active hand) for the beta band (see section 3.3.).

**Figure 5.**
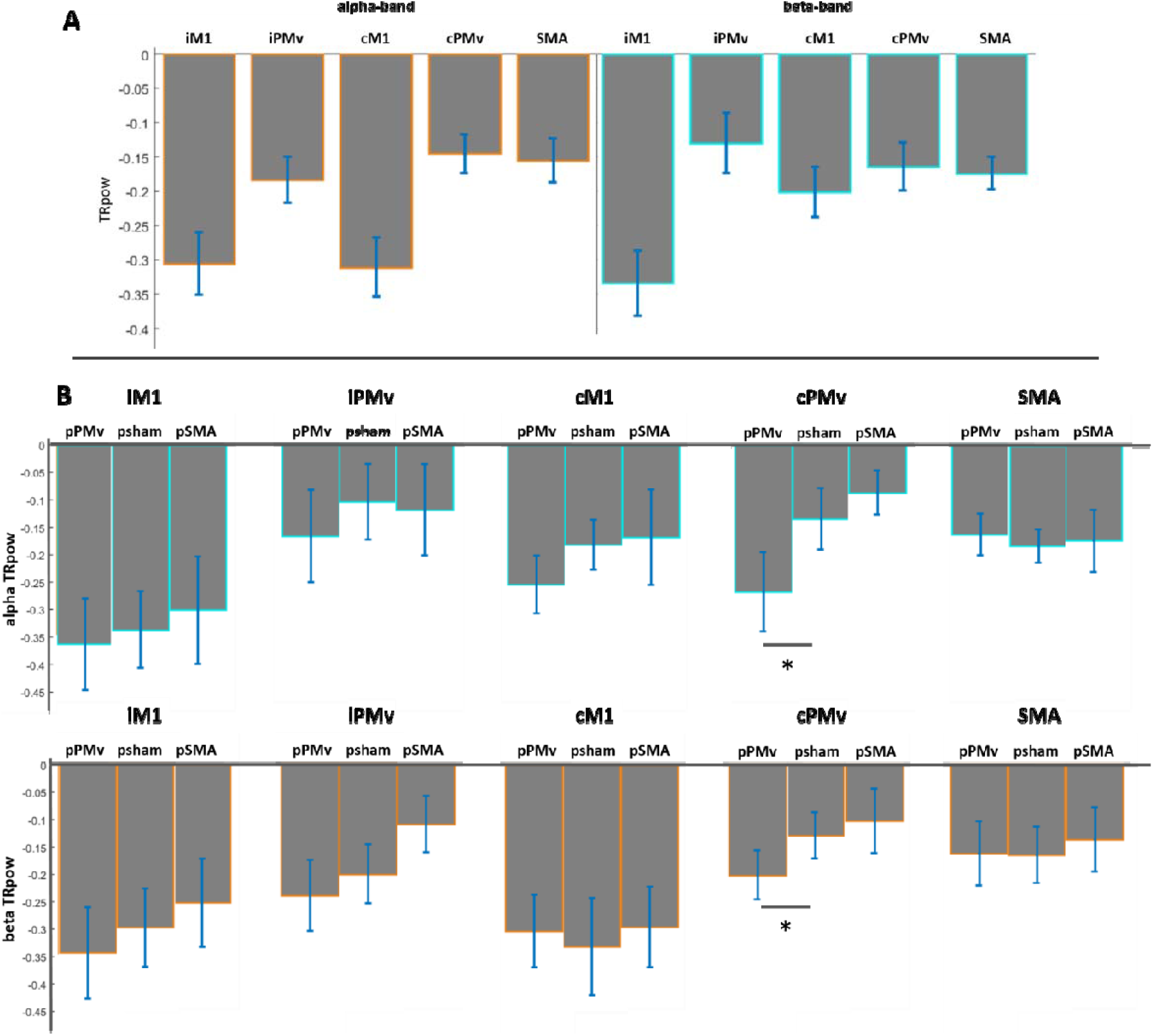
Task-related power changes. **A:** TRPD averaged over all experimental conditions, for alpha-band (left panel) and beta-band (right panel) at the source levels. TRPD (with standard errors) is plotted for iM1 (ipsilesional primary motor cortex, contralateral to the hand executing the grip task), cM1 (contralesional primary motor cortex), iPMv (ipsilesional ventral premotor cortex), cPMv (contralesional ventral premotor cortex), and SMA (supplementary motor area). **B:** TRPD for (i) alpha-band and beta-band and (ii) different stimulation sites at the source levels (pPMv = post stimulation of PMv, psham = post sham stimulation, pSMA = post stimulation of SMA). Reduction TRPow at contralesional PMv was significantly stronger (increased TRPD) both for the alpha (p = 0.0446) and beta-band (p = 0.0402) compared to sham stimulation.

As a first step, we assessed stimulation outlasting network modulation in form of local activity changes for each stimulation condition by comparing TRPD in the task block following each stimulation condition with the task block following sham stimulation. This revealed a significant reduction of task-related power, i.e., a significantly stronger TRPD at contralesional PMv for both alpha band (active stimulation, −0.20±0.16 vs. sham stimulation, - 0.13±0.17, p=0.0446) and beta band (active, −0.27±0.23vs. sham, −0.14±0.18, p=0.0402) after tACS over ipsilesional PMv (Fig. 5B). Order of stimulation was tested as a non-significant cofactor for these significant changes (Tab. S1 for details including FDR-corrected data (Benjamini & Hochberg, 1995), Supplementary Information).

### 3.3. Modulation of connectivity

We analyzed connectivity patterns (see section 2.5.2) between the nodes of the presented fronto-central motor network (see section 3.2.).

After 11Hz-PMv stimulation, the following changes compared to the period after sham stimulation were seen in the motor network during grip-task execution (all metrics task-related): reduced alpha coherence (active stimulation, −0.0386±0.0703 vs. sham stimulation, 0.0226±0.0694, p=0.0283) and phase-locking value (active, −0.0363±0.0581 vs. sham, 0.0036±0.0497, p=0.0097) between the ipsilesional M1 and contralesional PMv, reduced alpha coherence interhemispherically between the primary motor cortices (active, - 0.0514±0.0665 vs. sham, 0.0085±0.0888, p=0.0029), reduced imaginary alpha coherence between ipsilesional M1 and SMA (active, −0.0200±0.0641 vs. sham, 0.0153±0.0548, p=0.0462), reduced beta imaginary coherence between contralesional PMv and SMA (active, −0.0689±0.0628 vs. sham, −0.0058±0.0326, p=0.0101). While reduced task-related coupling values were the prominent observation, the alpha phase-locking value between contralesional M1 and SMA was increased (active, 0.0361±0.1209 vs. sham, - 0.0226±0.0923, p=0.0473). The order of stimulation conditions did not significantly influence the model fit.

After the 21Hz-SMA stimulation, the motor network during grip-task execution (all values task-related) showed a reduced beta coherence between ipsilesional M1 and contralesional PMv (active, −0.0367±0.0406 vs. sham, 0.0165±0.0737, p=0.0476). Beta-band imaginary coherence (active, −0.0224±0.0513 vs. sham, 0.0090±0.0259, p=0.0818) and beta-band phase-locking value (active, −0.0274±0.0356 vs. sham, 0.0115±0.0537, p=0.0577) were reduced revealing a statistical trend for this connection (iM1 to cPMv). The order of stimulation conditions did not significantly influence any of the model fits.

Figure 6 illustrates significant network modulations (see also Tab. S2 and S3 in the Supplementary Information, including FDR-corrected data).

**Figure 6.**
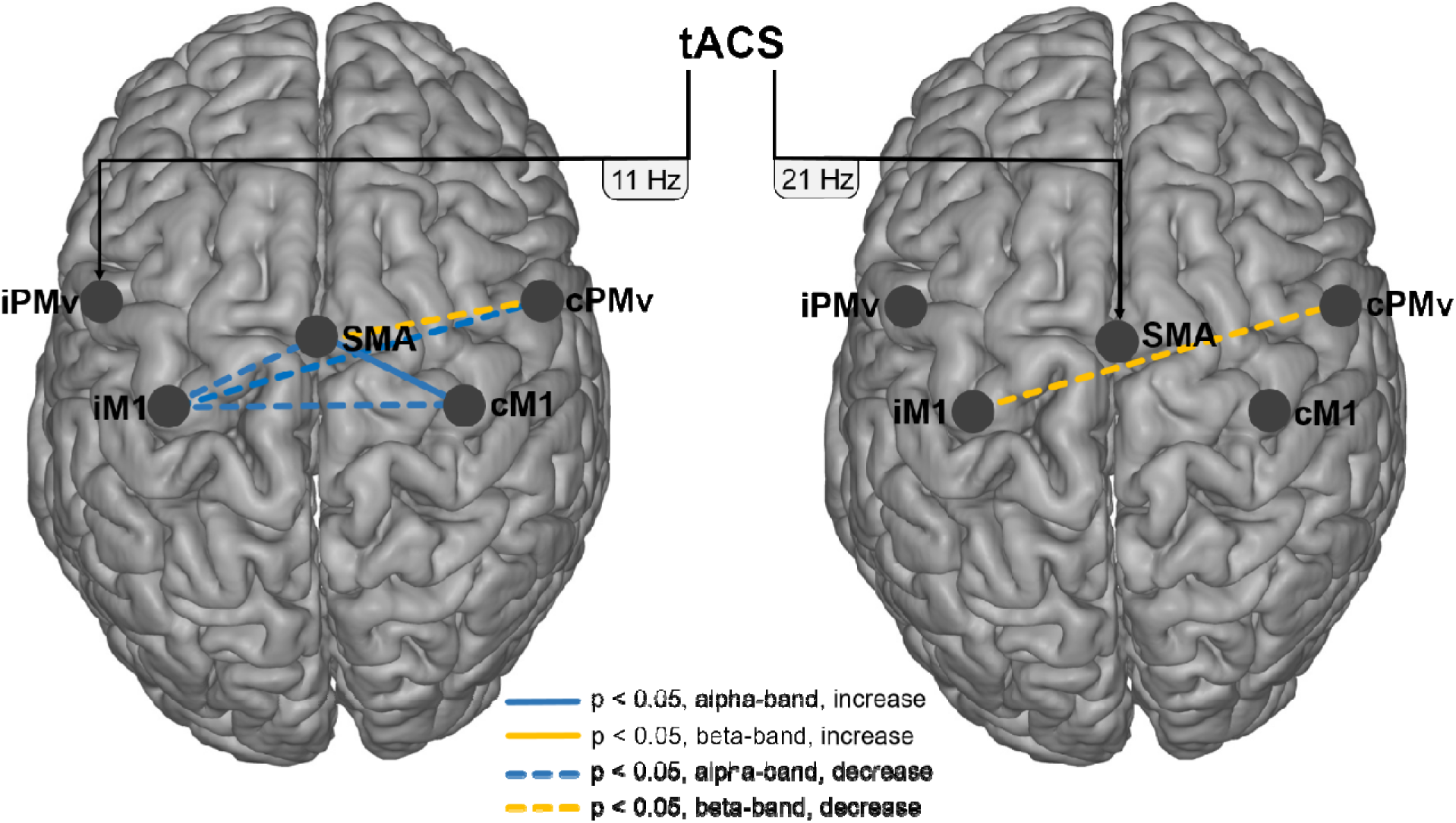
Modulated connections by 11Hz tACS over PMv (left panel) and 21Hz tACS over SMA (right panel). Connectivity was measured by coherence, phase-locking value, imaginary coherence and cross-frequency coupling (mutual information). M1 indicates primary motor cortex; PMv, ventral premotor cortex; SMA, supplementary motor area; i, ipsilesional; c, contralesional. Blue lines indicate alpha-band connectivity and yellow lines beta-band connectivity (Tab. S2 and S3). Dashed lines indicate decreases and solid lines increases. There were no significant changes in mutual information (Tab. S2 and S3).

### 3.4. Simulation of electric fields

We performed simulations of the applied electric fields elicited by the stimulation electrodes montages used in this study. To induce currents specifically, we used a very focal montage (Helfrich, Knepper, et al., 2014) with a moderate current of 1mA as described and illustrated in section 2.4. The simulation revealed a cortical electric field yield of about 0.06V/m for PMv stimulation and about 0.03V/m for SMA stimulation (Fig. 7).

**Figure 7.**
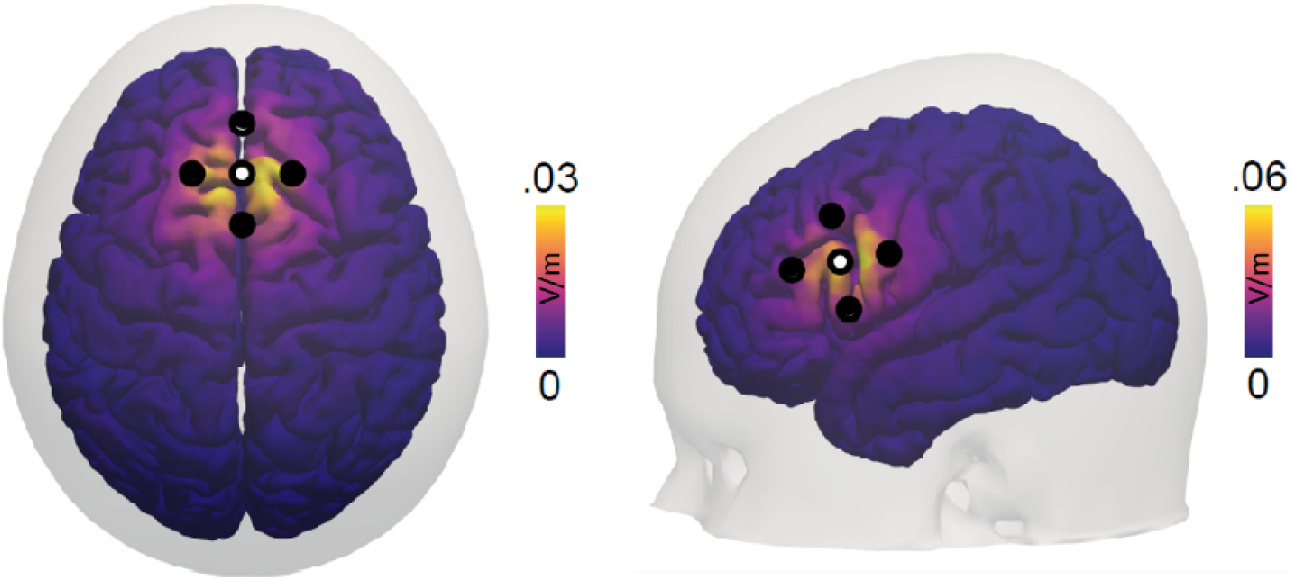
Simulation of cortical electric fields for stimulation of SMA (left panel) and PMv (right panel). Simulation revealed cortical electric fields of about 0.03V/m for SMA stimulation and 0.06V/m for PMv stimulation. We used a montage with four peripheral and one central electrode. Dots indicate electrode positions (white: inner electrodes, black: outer electrodes).

## 4. Discussion

In the present study, we investigated if task-related connectivity within a cortical motor network can be modified via low-intensity, focal tACS in chronic stroke patients. On the basis of previous work regarding changes of oscillatory activity after stroke in sensorimotor alpha and beta rhythms at core motor regions (Bönstrup et al., 2014, 2015, 2016) and recent evidence of tACS-effects on ongoing oscillations (Helfrich, Knepper, et al., 2014; Helfrich, Schneider, et al., 2014; Zaehle et al., 2010) we applied 11Hz tACS over ipsilesional PMv and 21Hz tACS over SMA with a focal electrode setup (Fig. 4). Given the broader aim of improving motor function by interfering with pathophysiologic network configuration during motor control, we applied stimulation and probed network architecture during the execution of a motor task (Bönstrup et al., 2015, 2016, 2018) (Fig. 4). With the scope of developing non-invasive brain stimulation as an “adjuvant” therapy in neurorehabilitation, our primary interest refers to stimulation effects outlasting the actual time of the tACS. We present the results of a wide exploration of network changes across frequencies and using various mutually validating and complementary measures of network connectivity.

### Safety

In our experiment on 16 chronic stroke patients, no adverse events (mild, moderate or serious) occurred. Overall, tACS application using a focal montage and with 1 mA peak-to-peak amplitude was safe and well tolerated in stroke patients. Patients reported minor skin sensations of mild tingling.

### tACS effects on regional neuronal activation

During the grip-task execution, TRPow at the sensorimotor alpha and beta rhythms decreased at all nodes within the motor network with a maximum at ipsilesional M1 (Fig. 5A), consistent with their role in sensorimotor engagement. There was a slight increase of alpha and beta-band TRPow decrease (TRPD) in the contralesional PMv after tACS applied to the ipsilesional PMv. Physiologically, stronger TRPD after tACS is considered to reflect increased local activation (Pfurtscheller & Lopes da Silva, 1999). Though being slight and not restricted to the stimulated frequency, the reduced TRPD at contralesional PMv might be mediated through transcallosal connections (Schulz et al., 2014). Since connectivity measures can be directly (amplitude correlations) or indirectly (signal-to-noise ratio) influenced by power amplitude, the relatively stable TRPow values across stimulation conditions are a good basis for the interpretation of the connectivity results (Bastos & Schoffelen, 2016).

### tACS effects on inter-regional connectivity

We conducted an exploratory analysis of coupling metrics before and after tACS by means of coherence, imaginary coherence, phase-locking value and cross-frequency coupling on the basis of mutual information. We found a consistent decrease of ipsilesional M1 and contralesional PMv connectivity for coherence and phase-locking value (section 3.3., Fig. 6, Tab S1/S2).

Anatomical and functional connections between ipsilesional PMv (stimulation target) and ipsilesional M1 are strong and of high relevance for motor control (Bönstrup et al., 2016; Rehme et al., 2011, 2012). Thus, stimulation of PMv may have indirectly impacted on the tightly connected M1 of the same hemisphere. Plasticity mechanisms after stroke involve a formation of new connections by secondary motor areas extending into the perilesional zone (Frost et al., 2019). The reduction of connectivity between ipsilesional M1 and contralesional M1/PMv could point to a more effective processing within the stroke-lesioned hemisphere after stimulation, requiring less support by an extended bilateral network. This would be in line with the concept of a reconstitution of lateralized, ipsilesional activity during movements in well recovered stroke patients (Rehme et al., 2011). Additionally, we found a significant reduction in contralesional PMv and SMA imaginary coherence in the beta band and a trend in the alpha band which could likewise be related to more efficient ipsilesional motor network processing after stimulation.

However, stronger TRPD at contralesional PMv, as discussed above, might rather point towards increased contralesional activation. Since TRPD reflects local phenomena, this finding is not necessarily at conflict with the finding of a reduction of connectivity between ipsilesional M1 and contralesional M1/PMv.

Beyond the observed tACS-induced reduction of interhemispheric coupling, there was an increase of the alpha-band phase-locking value between contralesional M1 and SMA and a decrease of alpha-band imaginary coherence between ipsilesional M1 and SMA. As the regulatory role of the SMA on M1 is well established (Ikeda et al., 1992), this finding might be interpreted in a similar way as the reduced interhemispheric coupling, that is, as an indicator that tACS may drive ipsilesional local oscillatory activity in M1, thereby suppressing input from secondary motor areas like SMA (Rehme et al., 2011) the contralateral M1 and PMv (Rehme & Grefkes, 2013).

Following beta-tACS to the SMA, task-related coherence between ipsilesional M1 and contralesional PMv was reduced in the beta band. In addition, there was a trend towards reduced alpha-band imaginary coherence and alpha-band phase locking value. In line with the findings after PMv stimulation, this reduction of connectivity between ipsilesional M1 and contralesional PMv, respectively, might point to a more active local processing within the stroke-lesioned hemisphere after stimulation.

For both PMv and SMA stimulation there were no significant effects on cross-frequency mutual information. In other words, the majority of effects seen were within the stimulated frequency band.

Taken together, we found weak tACS-effects on network architecture, by and large pointing to reduced interhemispheric and premotor-to-motor task-related connectivity. Potential reasons for the small effect sizes and sparse patterns of connectivity modulation are discussed in the next section.

### Limitations

One aspect of our study design that might be of relevance for the weak effects is the fact that stimulation was applied continuously during motor activity and resting periods between trials. Most tACS studies so far described after-effects following a stimulation of the resting brain (Schwab et al., 2018; Zaehle et al., 2010). TACS was shown to modulate brain activity via entrainment (Zaehle et al., 2010) and plasticity effects (particularly after-effects, (Vossen et al., 2015)). It is possible that the stimulation during periods of activation and rest elicited counter-effects on the local populational dynamics (Huang et al., 2005), thereby compromising the overall effect measured post stimulation.

Another aspect, which has to be addressed is the effective strength of tACS-induced fields which was potentially too weak. Simulations of stimulation setups applied in recent studies that showed an “offline” tACS effect on behavior or connectivity, revealed cortical field strengths at about 0,3V/m (Ahn et al., 2019), 0.37V/m (Schwab et al., 2018) or even 1V/m (Khatoun et al., 2018) in the stimulated area. Simulation of fields for our focal montage (Fig. 4) with a moderate current density of 1mA peak-to-peak revealed low fields compared to recent studies (Ahn et al., 2019; Khatoun et al., 2018; Schwab et al., 2018) of about 0.06V/m at the cortical surface in the area of PMv and 0.03V/m at the cortical surface in the area of SMA (Fig. 7). While this is an advantage for the focality of stimulation it might hamper the effectiveness, especially because our target regions were not located directly on the cortical convexity (Schulz et al., 2015, 2016). The weak effects of tACS in our study might be related to this low-intensity type of stimulation.

## 5. Conclusions

Taken together, we here present results of a study aimed to modulate the task-positive motor-network by tACS on PMv and SMA. In an exploratory analysis, we reveal a tACS-induced decreas of connectivity involving contralesional M1 and PMv, to some extent also SMA, potentially indicating stimulation-driven local ipsilesional neuronal activity after stimulation.

Major limitations result from applying tACS not at rest but during ongoing motor action and the choice of low stimulation intensity.

## 6. Acknowledgements

This research was supported by the German Research Foundation: SFB936 “Multi-site communication in the brain”, projects C1, C2, A1 and the German National Academy of Sciences, Leopoldina (LPDS 2016-01).

## 7. Conflicts of interest

We disclose financial or personal relationship with other people or organisations that could inappropriately influence this work.

## Declaration of interest

None.

