## Supplementary Information for "Effects of transcranial alternating-current stimulation to secondary motor areas on cortical oscillations in stroke patients"

1. **Modulation of TRPow**

Table S1 additionally shows P-values of linear mixed effects models for possible modulations of TRPow after tACS over PMv and SMA (section 3.2., Fig. 5B.).

**Table S1.** TRPow in detail.

|  | **p-values TRPow** | | | | |
| --- | --- | --- | --- | --- | --- |
|  | **iM1** | **iPMv** | **cM1** | **cPMv** | **SMA** |
| **post PMv stimulation** |  |  |  |  |  |
| alpha band | 0.3823 | 0.3360 | 0.6719 | **0.0446** | 0.9375 |
| *FDR-corrected* | *1.4549* | *1.4549* | *1.9177* | *0.5092* | *2.1406* |
| beta band | 0.6184 | 0.3285 | 0.1378 | **0.0402** | 0.6794 |
| *FDR-corrected* | *1.5513* | *1.2501* | *0.7866* | *0.4590* | *1.5513* |
| **post SMA stimulation** |  |  |  |  |  |
| alpha band | 0.4472 | 0.0926 | 0.6427 | 0.7032 | 0.4279 |
| *FDR-corrected* | *1.6056* | *1.0572* | *1.6056* | *1.6056* | *1.6056* |
| beta band | 0.5984 | 0.8377 | 0.8197 | 0.3379 | 0.8779 |
| *FDR-corrected* | *2.0045* | *2.0045* | *2.0045* | *2.0045* | *2.0045* |

**Table S1.** P-values of linear mixed effects models factor condition (post stimulation versus post sham) on TRPow during the motor task including FDR-corrected data. Both alpha-band TRPD and beta-band TRPD at contralesional PMv were significantly weaker after ipsilesional stimulation of PMv. Order of stimulation was tested as a non-significant cofactor for these significant changes. M1 indicates primary motor cortex; PMv, ventral premotor cortex; SMA, supplementary motor area; i, ipsilesional; c, contralesional.

2. **Modulation of connectivity**

Table S2 and table S3 additionally show p-values of linear mixed effects models for possible modulations of connectivity-measurements after tACS over PMv and SMA including FDR-corrected data (section 3.3., Fig. 6).

**Table S2**. P-values of connectivity measures post PMv stimulation.

| **p-values post PMv stimulation** | | | | | | | |
| --- | --- | --- | --- | --- | --- | --- | --- |
|  | **alpha band** | | | **beta band** | | | **cross-f** |
| **connections** | **coh** | **plv** | **i-coh** | **coh** | **plv** | **i-coh** | **MI** |
| **iM1-iPMv** | 0.1565 | 0.8642 | 0.2992 | 0.1249 | 0.5808 | 0.1740 | 0.0774 |
| **iM1-cM1** | **0.0029** | 0.1786 | 0.4297 | 0.8810 | 0.8810 | 0.9477 | 0.7393 |
| **iM1-cPMv** | **0.0283** | **0.0097** | 0.3801 | 0.9208 | 0.9343 | 0.3741 | 0.7663 |
| **iM1-SMA** | 0.3395 | 0.4126 | **0.0462** | 0.1438 | 0.2319 | 0.7768 | 0.1058 |
| **iPMv-cM1** | 0.3520 | 0.4645 | 0.1859 | 0.0921 | 0.6143 | 0.1113 | 0.0978 |
| **iPMv-cPMv** | 0.2890 | 0.2732 | 0.6422 | 0.3591 | 0.1749 | 0.9895 | 0.4177 |
| **iPMv-SMA** | 0.4000 | 0.9751 | 0.8376 | 0.5500 | 0.4390 | 0.3938 | 0.1896 |
| **cM1-cPMv** | 0.5399 | 0.6350 | 0.9042 | 0.2125 | 0.1643 | 0.8348 | 0.6405 |
| **cM1-SMA** | 0.0667 | **0.0473** | 0.6669 | 0.1987 | 0.7104 | 0.6341 | 0.1383 |
| **cPMv-SMA** | 0.7177 | 0.4206 | 0.0555 | 0.2366 | 0.4448 | **0.0101** | 0.9660 |
| **p-values post PMv stimulation - FDR corrected** | | | | | | | |
|  | **alpha band** | | | **beta band** | | | **cross-f** |
| **connections** | **coh** | **plv** | **i-coh** | **coh** | **plv** | **i-coh** | **MI** |
| **iM1-iPMv** | 1.1460 | 2.8125 | 2.0976 | 1.1550 | 2.5704 | 1.6988 | 1.0127 |
| **iM1-cM1** | 0.0849 | 1.7437 | 2.0976 | 2.6970 | 2.7365 | 2.8982 | 2.4939 |
| **iM1-cPMv** | 0.4144 | 0.2841 | 2.0976 | 2.6970 | 2.7365 | 2.3069 | 2.4939 |
| **iM1-SMA** | 1.4645 | 1.9436 | 0.8128 | 1.1550 | 2.2641 | 2.8982 | 1.0127 |
| **iPMv-cM1** | 1.4645 | 1.9436 | 1.8150 | 1.1550 | 2.5704 | 1.6300 | 1.0127 |
| **iPMv-cPMv** | 1.4645 | 1.9436 | 2.4417 | 1.5026 | 2.2641 | 2.8982 | 2.0391 |
| **iPMv-SMA** | 1.4645 | 2.8560 | 2.6484 | 2.0137 | 2.5704 | 2.3069 | 1.1107 |
| **cM1-cPMv** | 1.7571 | 2.3249 | 2.6484 | 1.1550 | 2.2641 | 2.8982 | 2.4939 |
| **cM1-SMA** | 0.6512 | 0.6927 | 2.4417 | 1.1550 | 2.6009 | 2.8982 | 1.0127 |
| **cPMv-SMA** | 2.1021 | 1.9436 | 0.8128 | 1.1550 | 2.5704 | 0.2958 | 2.8294 |

**Table S2.** **Top:** P-values of linear mixed effects models for factor condition (post stimulation vs. post sham) on coherence (coh), phase-locking value (plv) and imaginary coherence (i-coh) for alpha band and beta band after tACS over PMv. Moreover p-values for cross-frequency coupling (MI) are shown. M1 indicates primary motor cortex; PMv, ventral premotor cortex; SMA, supplementary motor area; i, ipsilesional; c, contralesional. Significant modulations: decreased alpha-band coherence iM1-cM1 (p = 0.0029), decreased alpha-band coherence iM1-cPMv (p = 0.0283), decreased alpha-band phase-locking value iM1-cPMv (p = 0.0097), decreased alpha-band imaginary coherence iM1-SMA (p = 0.0462), upregulated alpha-band phase-locking value cM1-SMA (p = 0.0473), decreased beta-band imaginary coherence cPMv-SMA (p = 0.0101). Order of stimulation was tested as a non-significant cofactor for these significant changes. **Bottom:** Corresponding FDR-corrected data. There was trend (p = 0.0849) for decreased alpha-band coherence between iM1-cM1 (p = 0.0029).

**Table S3**. P-values of connectivity measures post SMA stimulation.

| **p-values post SMA stimulation** | | | | | | | |
| --- | --- | --- | --- | --- | --- | --- | --- |
|  | **alpha band** | | | **beta band** | | | **cross-f** |
| **connections** | **coh** | **plv** | **i-coh** | **coh** | **plv** | **i-coh** | **MI** |
| **iM1-iPMv** | 0.3551 | 0.3285 | 0.6358 | 0.5382 | 0.5811 | 0.2737 | 0.0844 |
| **iM1-cM1** | 0.0947 | 0.5759 | 0.7106 | 0.1700 | 0.5512 | 0.1645 | 0.3347 |
| **iM1-cPMv** | 0.1389 | 0.2206 | 0.7558 | **0.0476** | 0.0577 | 0.0818 | 0.1932 |
| **iM1-SMA** | 0.5680 | 0.7981 | 0.1194 | 0.2791 | 0.3644 | 0.1696 | 0.6520 |
| **iPMv-cM1** | 0.5214 | 0.4435 | 0.7136 | 0.5117 | 0.7415 | 0.9647 | 0.7965 |
| **iPMv-cPMv** | 0.9049 | 0.4376 | 0.9539 | 0.2605 | 0.9522 | 0.3072 | 0.2069 |
| **iPMv-SMA** | 0.5303 | 0.9800 | 0.5474 | 0.6808 | 0.4148 | 0.3126 | 0.8016 |
| **cM1-cPMv** | 0.9052 | 0.7952 | 0.7518 | 0.3276 | 0.3250 | 0.3844 | 0.6888 |
| **cM1-SMA** | 0.1372 | 0.1603 | 0.5689 | 0.9029 | 0.3195 | 0.8436 | 0.4039 |
| **cPMv-SMA** | 0.3859 | 0.3751 | 0.1867 | 0.8250 | 0.7412 | 0.7878 | 0.7070 |
| **p-values post SMA stimulation - FDR correctred** | | | | | | | |
|  | **alpha -band** | | | **beta -band** | | | **cross-f** |
| **connections** | **coh** | **plv** | **i-coh** | **coh** | **plv** | **i-coh** | **MI** |
| **iM1-iPMv** | 2.0796 | 2.1650 | 2.4597 | 2.2520 | 2.4131 | 1.5260 | 2.0200 |
| **iM1-cM1** | 1.3561 | 2.4097 | 2.4597 | 1.9191 | 2.4131 | 1.5260 | 2.3479 |
| **iM1-cPMv** | 1.3561 | 2.1650 | 2.4597 | 1.3942 | 1.6900 | 1.5260 | 2.0200 |
| **iM1-SMA** | 2.0796 | 2.5973 | 2.4597 | 1.9191 | 2.4131 | 1.5260 | 2.3479 |
| **iPMv-cM1** | 2.0796 | 2.1650 | 2.4597 | 2.2520 | 2.4131 | 2.8256 | 2.3479 |
| **iPMv-cPMv** | 2.6513 | 2.1650 | 2.7939 | 1.9191 | 2.7890 | 1.5260 | 2.0200 |
| **iPMv-SMA** | 2.0796 | 2.8704 | 2.4597 | 2.4926 | 2.4131 | 1.5260 | 2.3479 |
| **cM1-cPMv** | 2.6513 | 2.5973 | 2.4597 | 1.9191 | 2.4131 | 1.6084 | 2.3479 |
| **cM1-SMA** | 1.3561 | 2.1650 | 2.4597 | 2.6446 | 2.4131 | 2.7454 | 2.3479 |
| **cPMv-SMA** | 2.0796 | 2.1650 | 2.4597 | 2.6446 | 2.4131 | 2.7454 | 2.3479 |

**Table S3. Top:** P-values of linear mixed effects models for factor condition (post stimulation vs. post sham) on coherence (coh), imaginary coherence (i-coh) and phase-locking value (plv) for alpha band and beta band after tACS over SMA. Moreover p-values for cross-frequency coupling (MI) are shown. M1 indicates primary motor cortex; PMv, ventral premotor cortex; SMA, supplementary motor area; i, ipsilesional; c, contralesional. Significant modulation: reduced beta-band coherence iM1-cPMv (p = 0.0476). Order of stimulation was tested as a non-significant cofactor for the significant change. **Bottom:** Corresponding FDR-corrected data.

3. **Behavioral performance**

In order to assess variable grip performance across stimulation blocks, we quantified the applied force during each stimulation and sham block. The average target grip force during each block, corresponding to 20% of maximal voluntary contraction in each patient, over all patients was 7.30 kg (± 2.54 SD). We computed the average applied grip force during the plateau-phase (1s to 9s) in the task-execution blocks following the stimulation. Following PMv stimulation, the applied force was 72.35 kg (± 25.25), following sham stimulation it was 71.43 kg (± 25.6) and SMA stimulation it was 71.88 kg (± 25.30). There were no significant differences between performances (p = 0.1040 for grip force after PMv stimulation versus grip force after sham stimulation, p = 0.1358 for grip force after SMA stimulation versus grip force after sham stimulation).

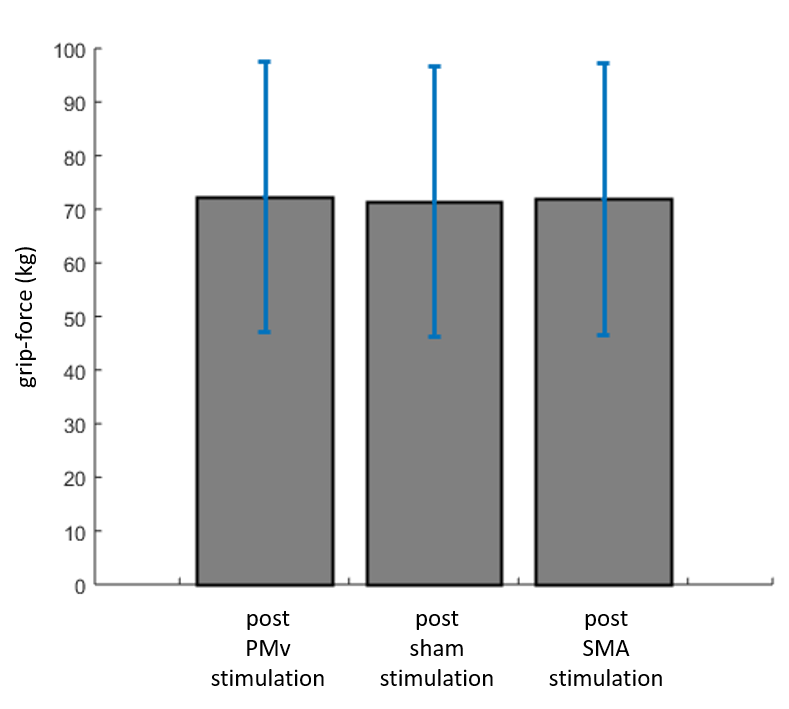

**Figure S1.** Force-production (kg) with standard deviation during gripping from 1s to 9s after the cue for the three conditions of stimulation. Patients performed blocks after PMv stimulation with 72.35 kg (± 25.25), blocks after sham stimulation with 71.43 kg (± 25.6) and blocks after SMA stimulation with 71.88 kg (± 25.30). There were no significant differences between both verum stimulation blocks and sham stimulation block.

4. **Resting-state analysis**

We additionally evaluated whether there was a change of local resting state-properties after tACS (Zaehle et al., 2010). However, since we designed our experimental procedures with a focus on task-related data, we did not include a resting-state epoch after the stimulation. Figure S2 exemplarily shows the limited possibilities of an analysis of a local change in phase-properties after stimulation. Only one non artifact-free episode of EEG per condition and patient would be able to be analyzed.

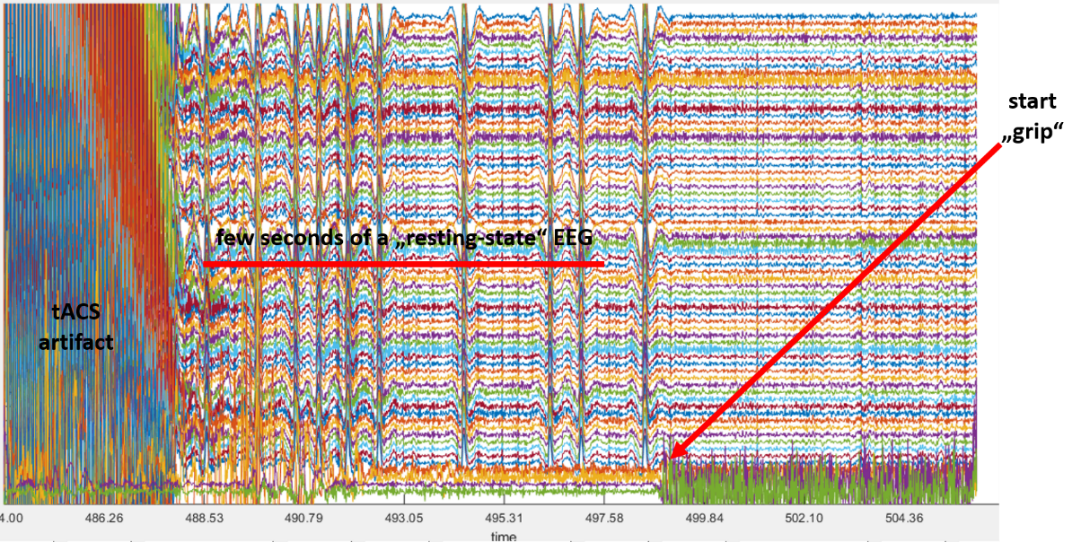

**Figure S2.** Possible epoch showing the full set of channels for a possible resting-state analysis. Due to the focus on task related activity, it was not possible to extract local resting state EEG after stimulation. Few seconds after the end of stimulation (artifact on the left) patients were instructed to start with gripping (EMG-activation “start-grip”).

5. **Outlook**

Recently, the role of parietal regions (anterior and posterior part of the intraparietal sulcus, called aIPS and cIPS) were shown to be relevant nodes in post-stroke plastic adaptations (Bönstrup et al., 2018; Schulz et al., 2016) and behaviorally relevant. For example, task-related alpha and beta-band coherence between aIPS and M1 was found to be increased in chronic stroke patients (Bönstrup et al., 2018). The strength of this connection also correlated with grip-force and performance in fine motor skills (Bönstrup et al., 2018). For future studies, parieto-frontal connections are promising targets for tACS network-modulation efforts, especially given the similarity of the target (parieto-frontal coherence) and stimulation method (alternating current at alpha or beta frequency).

Hence, we simulated an electric field using a less focal montage, targeting the parieto-frontal motor network including parietal areas relevant for visuo-motor integration (Bönstrup et al., 2018; Schulz et al., 2016). Moreover we simulated a higher current of 4mA leading to a spatially extended field with higher cortical voltages (Fig. S3).

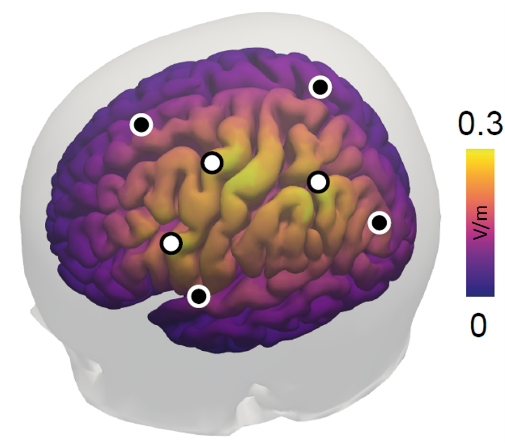

**Figure S3.** Simulation of cortical electric fields for an alternative montage yielding higher cortical currents. Simulation revealed cortical electric fields of about 0.3V/m. Dots indicate electrode positions (white: inner, cathodal electrodes, black: outer, anodal electrodes).

6. **References**

Bönstrup, M., Schulz, R., Schön, G., Cheng, B., Feldheim, J., Thomalla, G., & Gerloff, C. (2018). Parietofrontal network upregulation after motor stroke. *NeuroImage: Clinical*, *18*(March), 720–729. https://doi.org/10.1016/j.nicl.2018.03.006

Schulz, R., Buchholz, A., Frey, B. M., Bönstrup, M., Cheng, B., Thomalla, G., … Gerloff, C. (2016). Enhanced Effective Connectivity between Primary Motor Cortex and Intraparietal Sulcus in Well-Recovered Stroke Patients. *Stroke*, *47*(2), 482–489. https://doi.org/10.1161/STROKEAHA.115.011641

Zaehle, T., Rach, S., & Herrmann, C. S. (2010). Transcranial alternating current stimulation enhances individual alpha activity in human EEG. *PloS One*, *5*(11), e13766. https://doi.org/10.1371/journal.pone.0013766
